# A missense mutation in *TUBD1* is associated with high juvenile mortality in Braunvieh and Fleckvieh cattle

**DOI:** 10.1101/041921

**Authors:** Hermann Schwarzenbacher, Johann Burgstaller, Franz R. Seefried, Christine Wurmser, Monika Hilbe, Simone Jung, Christian Fuerst, Nora Dinhopl, Herbert Weissenböeck, Birgit Fuerst-Waltl, Marlies Dolezal, Reinhard Winkler, Oskar Grueter, Ulrich Bleul, Thomas Wittek, Ruedi Fries, Hubert Pausch

**Affiliations:** ZuchtData EDV Dienstleistungen GmbH, 1200 Vienna, Austria; Clinic for Ruminants, University of Veterinary Medicine Vienna, 1210 Vienna, Austria; Qualitas AG, 6300 Zug, Switzerland; Lehrstuhl fuer Tierzucht, Technische Universitaet Muenchen, 85354 Freising, Germany; Institute of Veterinary Pathology, Vetsuisse-Faculty, University Zurich, 8057 Zurich; Institute of Pathology and Forensic Veterinary Medicine, University of Veterinary Medicine, Vienna, Austria; Division of Livestock Sciences, University of Natural Resources and Life Sciences, Vienna, Austria; Platform Bioinformatics and Statistics, University of Veterinary Medicine, Vienna, Austria; ARGE Braunvieh, 6020 Innsbruck, Austria; Braunvieh Schweiz, 6300 Zug, Switzerland; Clinic of Reproductive Medicine, Department of Farm Animals, Vetsuisse-Faculty, University Zurich, 8057 Zurich, Switzerland

**Keywords:** Braunvieh haplotype 2, juvenile mortality, Tubulin delta 1, primary ciliary dyskinesia, ciliopathy, chronic respiratory disease

## Abstract

**Background:** Haplotypes with reduced or missing homozygosity may harbor deleterious alleles that compromise juvenile survival. A scan for homozygous haplotype deficiency revealed a short segment on bovine chromosome 19 (Braunvieh haplotype 2, BH2) that was associated with high juvenile mortality in Braunvieh cattle. However, the molecular genetic underpinnings and the pathophysiology of BH2 remain to be elucidated.

**Results:** The frequency of BH2 was 6.5 % in 8,446 Braunvieh animals from the national bovine genome databases. Both perinatal and juvenile mortality of BH2 homozygous calves were higher than the average in Braunvieh cattle resulting in a depletion of BH2 homozygous adult animals (P=9.3x10^−12^). The analysis of whole-genome sequence data from 54 Braunvieh animals uncovered a missense mutation in *TUBD1* (rs383232842, p.H210R) that was compatible with recessive inheritance of BH2. The availability of sequence data of 236 animals from diverse bovine populations revealed that the missense mutation also segregated at a low frequency (1.7 %) in the Fleckvieh breed. A validation study in 37,314 Fleckvieh animals confirmed high juvenile mortality of homozygous calves (P=2.2x10^−15^). Our findings show that the putative disease allele is located on an ancestral haplotype that segregates in Braunvieh and Fleckvieh cattle. To unravel the pathophysiology of BH2, six homozygous animals were examined at the animal clinic. Clinical and pathological findings revealed that homozygous calves suffered from chronic airway disease possibly resulting from defective cilia in the respiratory tract.

**Conclusions:** A missense mutation in *TUBD1* is associated with high perinatal and juvenile mortality in Braunvieh and Fleckvieh cattle. The mutation is located on a common haplotype likely originating from an ancient ancestor of Braunvieh and Fleckvieh cattle. Our findings demonstrate for the first time that deleterious alleles may segregate across closed cattle breeds without recent admixture. Homozygous calves suffer from chronic airway disease resulting in poor growth performance and high juvenile mortality. The respiratory manifestations resemble key features of diseases resulting from impaired function of airway cilia.

## Background

Reducing calf mortality is an important objective in cattle breeding populations both for animal welfare and economic reasons. Bovine calf losses including late abortions, stillbirths and diseases during rearing may be in excess of 10 % [1–4]. Most rearing losses are attributable to infectious diarrheal and respiratory diseases [5]. While a number of genetic variants predisposing to high perinatal mortality have been identified in bovine populations (e.g., [6, 7]), the mapping of loci affecting disease susceptibility and rearing success is difficult because of their low heritability [8, 9].

Case-control association testing using genome-wide marker data facilitated the discovery of causal variants for monogenic disorders that result in high juvenile mortality [10–16]. Although the pathophysiology of such conditions is heterogeneous, pertinently affected calves may be apathetic, without vigor, retarded in growth or highly susceptible to infectious disease. Affected calves may be born without typical signs of disease but develop clinical features shortly after birth [10, 11, 14].

The availability of large-scale genotype data enables the identification of haplotypes with homozygosity depletion [14, 17, 18]. Haplotypes with reduced or missing homozygosity in adult animals are likely to harbor recessive alleles that are associated with an increased pre-, perior postnatal mortality. Genome-wide scans for homozygous haplotype deficiency in North American [17] and Austrian Braunvieh [19] populations uncovered a short segment on bovine chromosome 19 that was associated with high postnatal calf losses (OMIA 001939-9913). The segment associated with calf mortality was denominated 424.49 and BTA19-1 in the North American [17] and Austrian Braunvieh [19] populations, respectively. Since 2013, the associated genomic segment is consistently referred to as BH2 with BH being an abbreviation for Braunvieh haplotype [20].

In the present study, we exploit comprehensive genotyping and sequencing data to detect a missense mutation in *TUBD1* that is most likely causal for the high juvenile mortality of BH2 homozygous calves. We validate the missense mutation in an independent cattle population and provide evidence that homozygous calves suffer from chronic respiratory disease.

## Results

### BH2 compromises juvenile survival in Braunvieh cattle

We exploited array-derived genotypes of 8,446 Braunvieh animals from the national bovine genome databases to scan for homozygosity depletion on chromosome 19. Thirty-two haplotypes in strong linkage disequilibrium that were located between 3.67 Mb and 13.38 Mb on BTA19 showed a significant depletion of homozygous animals (P<1x10^−4^). The length of the haplotypes with homozygosity depletion ranged from 0.54 to 6.33 Mb. Harmful effects on fertility and calf survival were analyzed for all haplotypes with homozygosity depletion using logistic regression analyses with insemination and rearing success as response variables. None of the haplotypes tested was associated with insemination success (P>0.01) ruling out homozygous haplotype deficiency to result from early embryonic losses. However, all haplotypes were associated with high juvenile mortality (P<3.2 x 10^−9^). After combining the P values from homozygosity depletion and calf survival analyses, the most significantly associated haplotype was located within a 1.14 Mb interval between 10,694,269 bp and 11,833,182 bp on BTA19 (**Figure 1a**). From now on, we refer to this haplotype as BH2. The frequency of BH2 was 6.5%. Only one animal was homozygous although 41 were expected (P=9.3x10^−12^). The analysis of 32,100 calving records revealed that the first-year mortality of descendants from BH2 risk-matings was increased by 4.6 *%* compared to non-risk matings (P=1.8x10^−28^)(**Table 1**).

**Figure 1:**
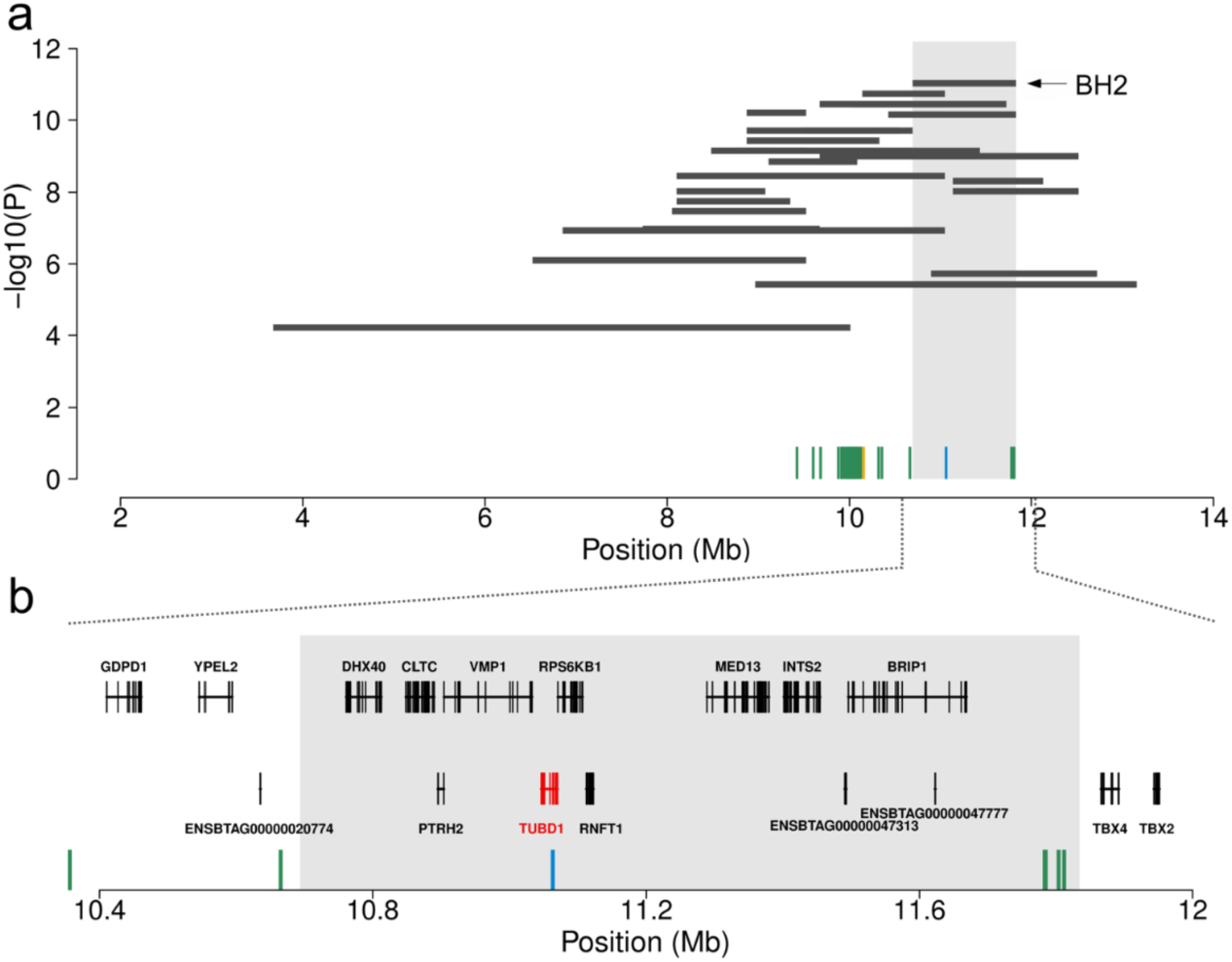
Schematic representation of the proximal region of bovine chromosome 19 encompassing BH2. Association of 32 haplotypes with homozygosity depletion and postnatal calf mortality (a). Each horizontal bar represents a haplotype with homozygosity depletion. The grey shaded area highlights the most significantly associated haplotype (BH2). Green, orange and blue vertical bars represent 50 non-coding variants, one synonymous variant in *PPM1E* and one missense variant in *TUBD1,* respectively. Genes located within the BH2 interval (b). The grey shaded area highlights the position of BH2. Green and blue vertical bars represent six non-coding variants and the missense variant in the *TUBD1* gene.

**Table 1:**
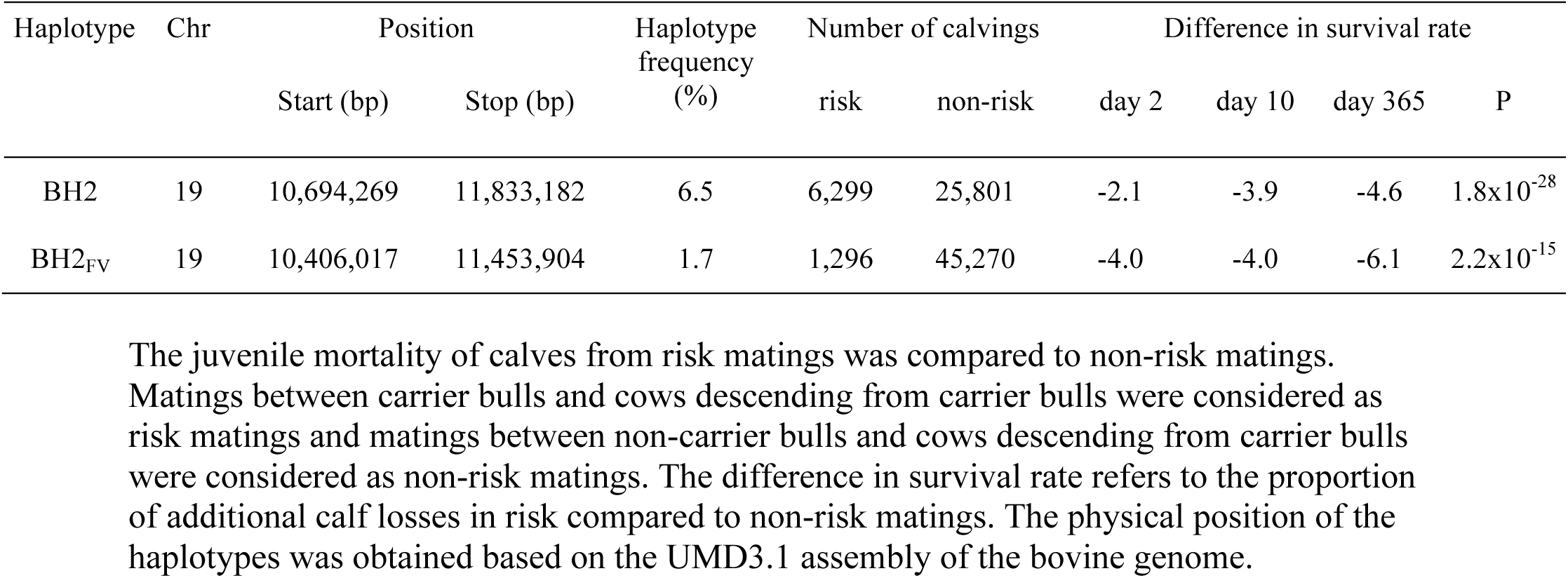
Two haplotypes with reduced homozygosity in Braunvieh (BH2) and Fleckvieh (BH2FV) cattle.

Two male Braunvieh calves homozygous for BH2 were detected in the Swiss bovine genome database. According to the possessing farmers, both calves were underweight at birth. One calf died for an unknown cause at 72 days of age and was not available for phenotyping. The second calf (BH2_hom_) was referred to the animal clinic at 66 days of age. During a hospitalization period of 95 days, BH2_hom_ suffered repeatedly from bronchopneumonia. At 161 days of age, BH2_hom_ was euthanized because of suddenly occurring severe dyspnea.

The juvenile mortality of calves from risk matings was compared to non-risk matings. Matings between carrier bulls and cows descending from carrier bulls were considered as risk matings and matings between non-carrier bulls and cows descending from carrier bulls were considered as non-risk matings. The difference in survival rate refers to the proportion of additional calf losses in risk compared to non-risk matings. The physical position of the haplotypes was obtained based on the UMD3.1 assembly of the bovine genome.

### A missense mutation in *TUBD1* is compatible with recessive inheritance of BH2

To identify the causal mutation for the high mortality of BH2 homozygous calves, we analyzed re-sequencing data of BH2_hom_, five heterozygous BH2 carriers and 48 control animals of the Braunvieh population. The average genome coverage in BH2_hom_, five heterozygous carriers and 48 control animals, respectively, was 17.36, 12.08±1.39 and 11.46±1.54 fold. Multi-sample variant calling in 54 sequenced animals yielded genotypes for 99,797 single nucleotide and short insertion and deletion polymorphisms and 10,497 structural variants located within the 9.71 Mb interval (from 3.67 Mb to 13.38 Mb) with homozygosity depletion encompassing BH2. These 110,294 polymorphic sites were filtered for variants that were compatible with recessive inheritance of BH2 that is homozygous for the alternate allele in BH2_hom_, heterozygous in BH2 carriers and homozygous for the reference allele in control animals. This filtering revealed 52 variants in LD with BH2 that were located between 9,428,803 bp and 11,811,557 bp on BTA19 (**Additional File 1)**: fifty variants were located in non-coding regions, one variant (rs479748045) was a synonymous mutation in the *PPM1E* (*protein phosphatase, Mg2+/Mn2+ dependent 1E*) gene and one variant (rs383232842) was a missense mutation in the *TUBD1* (*tubulin delta 1*) gene (**Additional File 2**). Of the 52 variants in LD with BH2 four non-coding variants and the missense mutation in *TUBD1* were located within the 1.14 Mb interval (BH2) that showed the strongest association with homozygosity depletion and postnatal calf mortality (**Figure 1b**). The four non-coding variants that were located within the BH2 interval were more than 50 kb away from coding sequences. Moreover, two of them (rs385391620 at 11,782,742 bp and rs386039720 at 11,803,361 bp) were found to occur in homozygous state among 1682 animals from various bovine breeds that had been sequenced for the 1000 bull genomes project (**Additional File 1**) [21] and are thus less likely to be causal for the high postnatal mortality associated with homozygosity for BH2. In conclusion, the rs383232842 mutation in the *TUBD1* coding region and two non-coding variants were considered as candidate causal variants underlying BH2 (**Additional File 3**). The rs383232842 C-allele causes a substitution of a histidine by an arginine at a conserved position in tubulin delta 1 (ENSBTAP00000001700.5:p.H210R) (**Additional File 4**). The amino acid substitution is predicted to be damaging to protein function (SIFT-score: 0.03, Polyphen-score: 0.23). We obtained genotypes of the rs383232842 polymorphism in 661 adult Braunvieh animals using a KASP genotyping assay; the mutation was in high linkage disequilibrium (LD) with BH2 (r^2^=0.98) and none of 661 genotyped animals was homozygous for the C-allele (**Table 2**). Genotyping results differed from the haplotype-based BH2 states for five (out of 661) animals possibly because of haplotype recombination or phasing errors. The rs383232842 polymorphism was in LD with the two non-coding variants (**Additional File 1** & **Additional File 5**).

**Table 2:** Genotypes for a missense mutation in TUBD1 in 661 adult Braunvieh cattle.

| BH2 haplotype status | N | Genotype at rs383232842 |  |  |
| --- | --- | --- | --- | --- |
|  |  | T/T | T/C | C/C |
| Non-carrier | 221 | 220 | 1 | - |
| Carrier | 440 | 4 | 436 | - |

Genotypes for the rs383232842 polymorphism were obtained in 661 Braunvieh animals using KASP genotyping assays. The haplotype status (non-carrier, carrier) of the animals was determined using phased genotypes obtained from the Illumina BovineSNP50 array. Note that the genotyped animals are not representative for the entire Braunvieh population because we preferentially selected heterozygous BH2 haplotype carriers for genotyping.

### The BH2 haplotype affects rearing success in another cattle breed

The analysis of whole-genome sequencing data from 236 animals from six bovine breeds other than Braunvieh revealed that rs383232842 was heterozygous in seven out of 149 Fleckvieh animals (**Additional File 1** & **Additional File 3**). To obtain genotypes for rs383232842 in a representative sample of the Fleckvieh population, we genotyped 3807 randomly selected adult animals using a KASP genotyping assay. Among these, 142 were heterozygous and none was homozygous for the C-allele corresponding to a C-allele frequency of 1.86%. Of 3807 animals that had been genotyped at rs383232842, 1974 were also genotyped with the Illumina BovineHD Bead chip comprising 777,962 SNPs. The availability of dense marker data enabled us to detect an 1.05 Mb haplotype (BH2_FV_; from 10,406,017 bp to 11,453,904 bp) that was specific for the rs383232842 C-allele: ninety-two animals that carried the C-allele were heterozygous carriers of BH2_FV_, whereas 1882 animals that were homozygous for the reference allele did not carry BH2_FV_. In 37,314 Fleckvieh animals of the national bovine genome databases that had been genotyped with the Illumina BovineSNP50 Bead chip, the frequency of BH2_FV_ was 1.68%. Only four animals were homozygous for BH2_FV_ although ten were expected (P=0.09). The analysis of 46,566 calving records revealed that the first-year mortality of descendants from BH2_FV_ risk matings was increased by 6.1 % compared to progeny from non-risk matings (P=2.2x10^−15^) (**Table 1**).

### A common haplotype with the rs383232842 C-allele segregates in Braunvieh and Fleckvieh cattle

BH2 encompasses a 1.14 Mb segment from 10,694,269 bp to 11,833,182 bp. It is defined by eleven SNPs of the Illumina BovineSNP50 Bead chip. BH2_FV_, defined by 249 SNPs of the Illumina BovineHD Bead chip, is located between 10,406,017 bp and 11,453,904 bp. The BH2 alleles are a subset of the BH2_FV_ alleles within a 741 kb segment between 10,694,269 bp and 11,435,269 bp (**Additional File 6**). Since both BH2 and BH2_FV_ contain the rs383232842 C-allele, a common origin of the two haplotypes is most likely. The analysis of pedigree records enabled us to track BH2 and BH2_FV_ back to the Braunvieh bull Rancho Rustic My Design (birth year 1963) and the Fleckvieh bull Polzer (birth year 1959), respectively. However, we were not able to identify common ancestors in their pedigrees possibly because of missing pedigree information from very distantly related relatives. Cluster analyses using genome-wide marker data of 442 Fleckvieh and 280 Braunvieh animals born between 1970 and 2010 revealed no evidence for an exchange of genetic material between both breeds (**Additional File 7**).

### BH2 segregates in the Holstein breed, but does not contain the rs383232842 C-allele

To test if the common BH2 / BH2_FV_ haplotype segment segregated in another dairy breed, we analyzed haplotypes of 8840 adult Holstein animals that had been genotyped with the Illumina BovineSNP50 Bead chip. We identified 917 heterozygotes and 30 homozygotes for BH2 among the genotyped Holstein animals. The observed haplotype distribution did not deviate from the Hardy-Weinberg equilibrium (P=0.54). We additionally genotyped the rs383232842 polymorphism in 503 Holstein bulls using a KASP genotyping assay. All of them including 74 heterozygotes and two homozygotes for BH2 were homozygous for the rs383232842 reference allele (**Additional File 6**). Another 311 Holstein animals that had been sequenced within the 1000 bull genomes project [21] were homozygous for the reference allele indicating that the rs383232842 C-allele does not segregate in this breed.

The non-reference alleles at two non-coding variants located within the BH2 interval were only observed at very low frequency in Holstein cattle (**Additional File 1 & Additional File 5**).

### Identification of BH2 homozygous calves

To unravel the pathophysiology associated with BH2, we initiated a monitoring project in the Austrian and Swiss Braunvieh populations. Breeding consultants collected ear tissue samples of calves descending from BH2 risk matings immediately after birth for genetic investigations. Of 117 genotyped calves, twelve were homozygous for the rs383232842 Callele (**Table 3**). Among those, eight were stillborn or died shortly after birth. All stillborn homozygotes were underweight at 25 to 30 kg and appeared underdeveloped although gestation length was normal (**Additional file 8**). Apart from low body weight, necropsy revealed no signs of disease. Four live born homozygous calves (BV1-BV4) were referred to the animal clinic.

**Table 3:** Sampling of calves from BH2 risk matings.

| Genotype at<br>rs383232842 | Number of<br>calves | Stillborn<br>calves | Died within<br>10 days | Died within<br>30 days |
| --- | --- | --- | --- | --- |
| TT | 45 | 1 | 0 | 1 |
| CT | 60 | 4 | 1 | 0 |
| CC | 12 | 5 | 0 | 3 |

Genotypes of 117 calves descending from BH2 risk matings. Note the high perinatal mortality (8/12) of calves homozygous for the rs383232842 C-allele.

During the sampling of BH2 homozygous calves, we encountered a male Fleckvieh calf (FV1) with low birth weight and postnatal growth restriction that descended from a BH2_FV_ carrier bull. At 40 days of age, its weight was 28.5 kg, which is only a third compared to healthy Fleckvieh calves of the same age. FV1 was tested negatively for two recessive mutations that are known to cause retarded growth [14, 22]. Another 16-months old Braunvieh bull (BV5) with poor growth performance and a history of recurrent respiratory disease was reported by a farmer. Inspection of the bull’s pedigree revealed BH2 carriers among its paternal and maternal ancestors. Sanger sequencing of DNA samples from FV1 and BV5 confirmed that both animals were homozygous for the rs383232842 C-allele. Both animals were referred to the animal clinic.

### Homozygous animals suffer from chronic airway disease

At admission, six homozygous animals were emaciated and their heads appeared elongated (**Figure 2, Additional file 9**). Initial examination revealed aberrant breathing sounds (*i.e.,* enforced vesicular breathing, rhonchus, coughing), tachypnea, tachycardia and excessive mucous exudation from the nostrils (**Table 4**). The analysis of blood parameters revealed iron deficiency and increased monocyte counts in some animals possibly indicating ongoing response to infectious disease (**Additional file 10**). The clinical features led to the diagnosis of bronchopneumonia and the animals were treated accordingly. Although the medication alleviated the clinical symptoms, respiratory disease recurred repeatedly. Due to the steadily declining health condition with no prospect for improvement, all animals were euthanized and subjected to necropsy.

**Figure 2:**
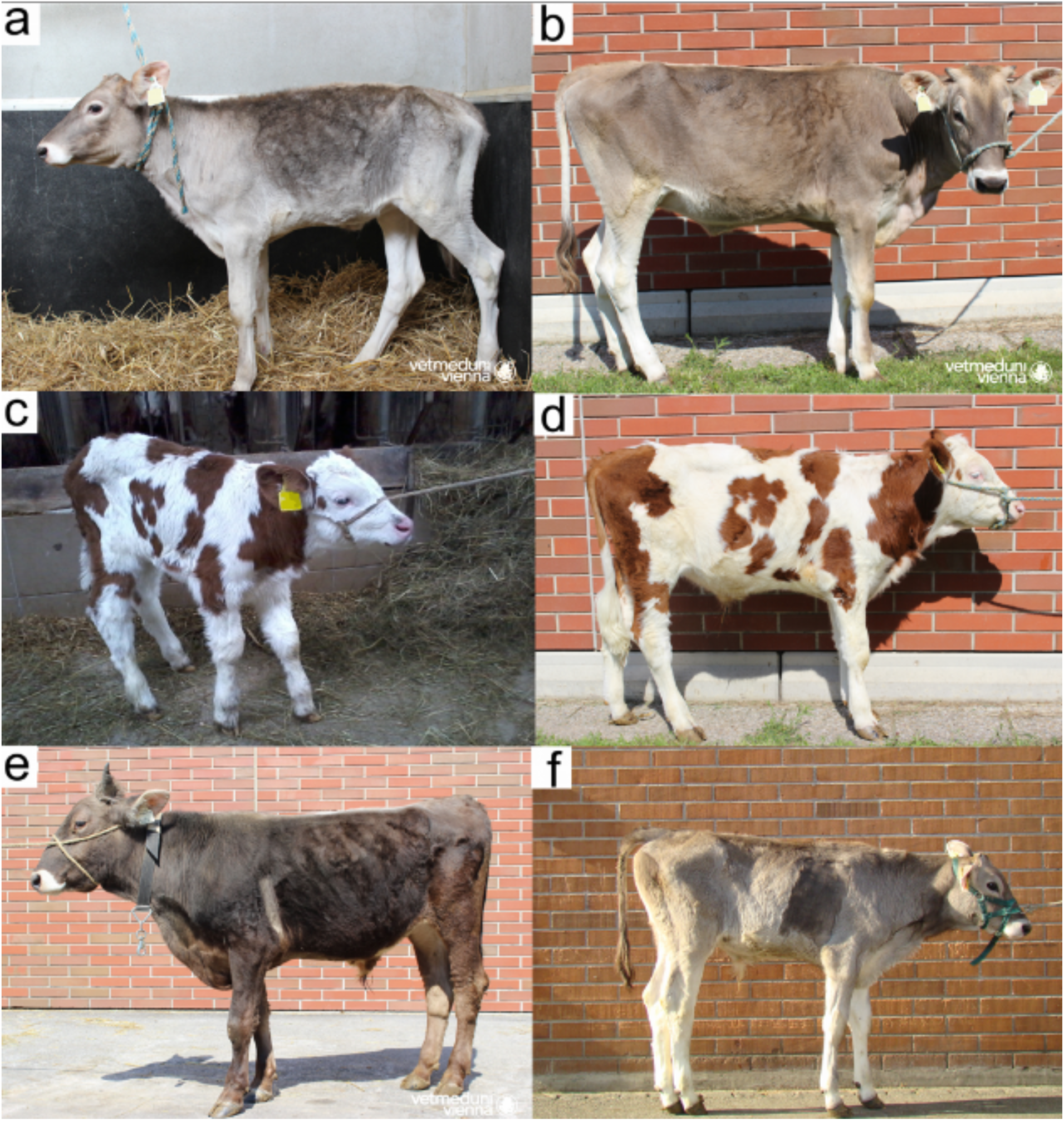
Phenotypic manifestation of homozygosity for BH2 in four animals. Pictures of BV2 (a, b), FV1 (c, d), BV5 (e) and BV4 (f) were taken at the time of admission in the animal clinic (a, c) and shortly before euthanasia (b, d, e, f), respectively. A detailed description of the disease symptoms is available in Table 4.

**Table 4:** Characteristics of six hospitalized animals.

| Parameter | BV1 | BV2 | FV1 | BV3 | BV4 | BV5 |
| --- | --- | --- | --- | --- | --- | --- |
| Breed | Braunvieh | Braunvieh | Fleckvieh | Braunvieh | Braunvieh | Braunvieh |
| Sex | male | female | male | male | male | male |
| Age at admission (days) | 41 | 44 | 75 | 13 | 1 | 485 |
| Weight at admission (kg) | 58 | 59 | 71 | 31 | 33 | 227 |
| Pulse (heartbeats per minute) | 100 | 104 | 100 | 140 | 138 | 60 |
| Body temperature (°C) | 38.3 | 39.9 | 39.2 | 38.6 | 38.3 | 39.1 |
| Respiratory rate (breaths per minute) | 40 | 48 | 40 | 40 | 54 | 60 |
| BCS (score) | 2.5 | 2 | 2.5 | 2 | 2 | 1.5 |
| Auscultation lungs | + vesicular | ++ vesicular | ++ vesicular | +++ vesicular | rhonchus | +++ vesicular<br>bronchial |
| Hospitalization period (days) | 109 | 164 | 111 | 19 | 157 | 17 |
| Age at euthanasia (days) | 150 | 208 | 186 | 32 | 158 | 502 |
| Weight at euthanasia (kg) | 168 | 151.5 | 106.5 | 31 | 134 | 258 |

Parameters from the initial examination of five Braunvieh (BV1-BV5) and one Fleckvieh (FV1) animal homozygous for the rs383232842 C-allele.

At necropsy, the animals were underweight and emaciated. Macroscopic abnormalities of the respiratory tract such as mucopurulent rhinitis, tracheobronchitis and lung lesions became evident in all animals (**Additional file 11**). Histological sections of tissue samples from the upper and lower respiratory tract revealed hyperplastic ciliated epithelia with intraepithelial neutrophils and bronchus-associated lymphoid tissue (BALT) hyperplasia, respectively, accompanied by bronchiolitis and bronchiectasis (**Additional file 12**). The lumen of bronchi and bronchioles were obstructed by mucopurulent exudate and the surrounding lung tissue was atelectatic and showed minimal intra-alveolar infiltration of inflammatory cells. Transmission electron microscopic (TEM) data of ciliated epithelial tissue were available from the upper and lower respiratory tract of four animals (FV1, BV1, BV2, BV4). Analyses of TEM sections revealed defective microtubule organization in 20-30 % of ciliary cross-sections including transposition faults, microtubular disorganization, absence of outer and central microtubule pair and aberrations of the outer and inner dynein arms (**Figure 3**).

**Figure 3:**
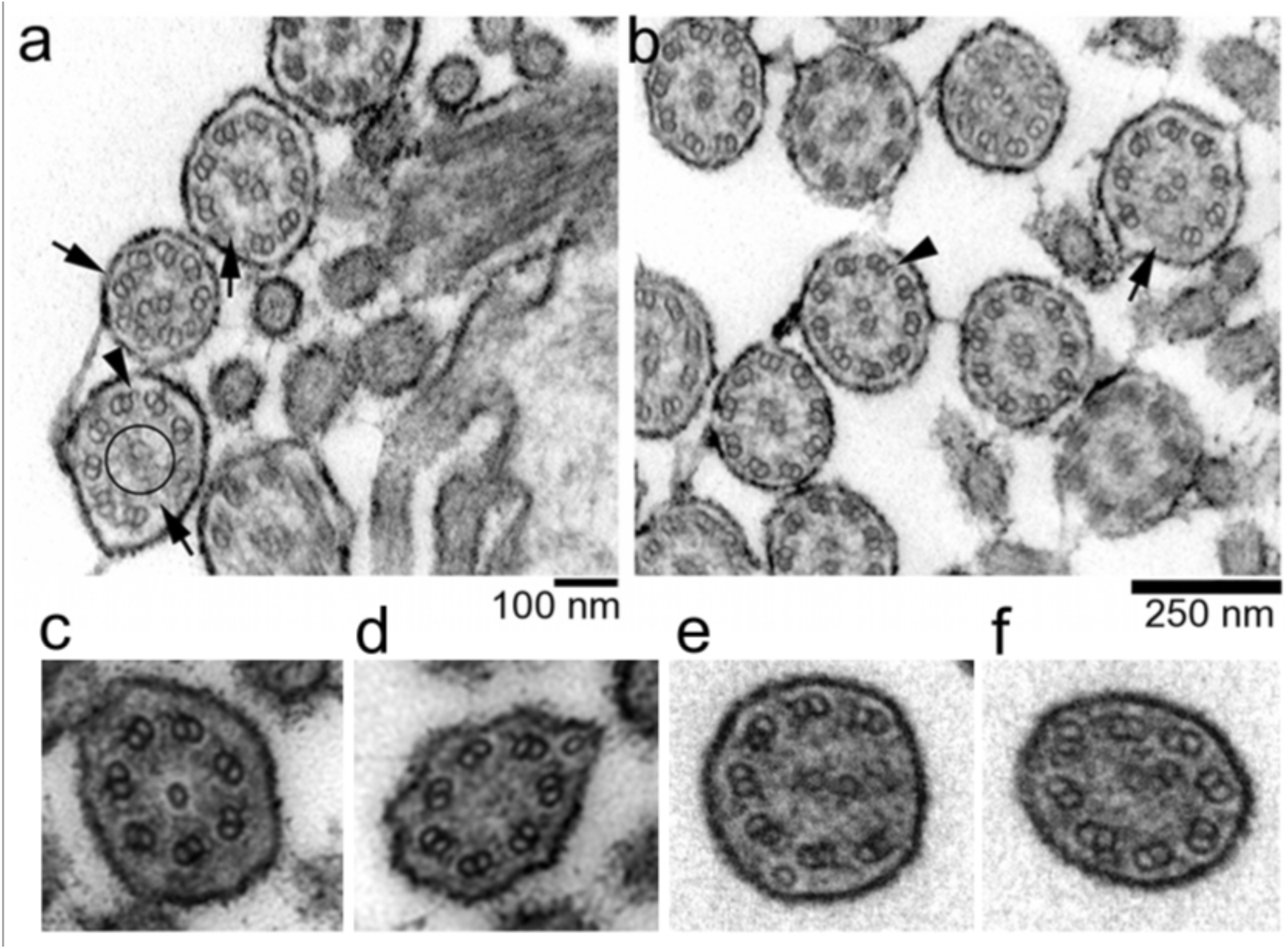
Transmission electron microscopy of respiratory cilia of homozygous animals. Motile cilia are characterized by a typical “9x2+2” architecture, *i.e.,* nine outer microtubule doublets surrounding a central pair of microtubule. The TEM sections of respiratory cilia of bronchi from BV1 (a) and FV1 (b) revealed multiple ultrastructural defects including transposition defects of microtubuli and microtubular disorganization (arrows), absence of a central microtubule pair (circle), and loss of inner and/or outer dynein arms (arrowheads). Magnification of cilia with typical ultrastructural defects found in the respiratory tract of BV5 (c,d) and BV4 (e,f).

### Homozygosity for BH2 may be incompletely penetrant

Breeding consultants reported another three Braunvieh animals that were homozygous for BH2. These homozygous animals were detected during routine genomic evaluation using genotypes of the Illumina BovineSNP50 Bead chip. Sanger sequencing of DNA samples confirmed homozygosity for the rs383232842 C-allele in all animals. While one young bull (BV6) was healthy and normally developed at the age of 370 days, two animals (BV7, BV8) were retarded in growth and had a history of recurrent respiratory disease (**Additional file 13**). The three homozygous animals were also homozygous for the alternate allele at two non-coding variants in LD with rs383232842 (**Additional File 5**).

## Discussion

We show that the proximal region of bovine chromosome 19 is associated with high peri-and postnatal calf losses in Braunvieh cattle, corroborating previous reports of a harmful haplotype (Braunvieh haplotype 2, BH2) in European and North-American Braunvieh populations [17, 19]. The joint analysis of homozygosity depletion and insemination and rearing data revealed a 1.14 Mb haplotype located between 10,694,269 bp and 11,833,182 bp on BTA19 as the most likely interval harboring the causal mutation, which agrees with previous findings [17, 19]. The first-year mortality of calves descending from BH2 risk matings is less than expected for a lethal recessive disorder, indicating that some BH2 homozygous calves may reach adulthood. Despite its detrimental effect on juvenile survival, the frequency of BH2 is high in the European Braunvieh populations (6.5 %). Based on the retrospective analysis of inadvertent BH2 risk-matings that happened in the Austrian Braunvieh population (N=50,000 calvings per year) between 1990 and 2013, we estimated that between 11 and 393 BH2 homozygous animals were born annually in the past 25 years. Deleterious alleles may reach high frequency in cattle populations because of the widespread use of unnoticed carrier bulls in artificial insemination and pleiotropic effects on desirable traits [23–25]. So far, there is no evidence that BH2 has desirable effects on important breeding objectives. Several harmful conditions other than BH2 segregate in the Braunvieh populations [17, 26–29]. Pinpointing the causal mutation allows for the implementation of efficient genome-based mating programs to avoid the inadvertent mating of carrier animals while maintaining genetic diversity and high rates of genetic gain. A missense mutation (rs383232842, p.H210R) in *TUBD1* was found to be compatible with recessive inheritance of BH2. Affected, carrier and non-carrier animals were homozygous for the alternate allele, heterozygous and homozygous, respectively, for the reference allele at rs383232842. Although BH2 is in high LD with rs383232842, the identification of carrier animals is less reliable with haplotype information. Five out of 661 animals were misclassified based on the haplotype test indicating both lower specificity and sensitivity of haplotype-based classification of animals into carriers and non-carriers which agrees with previous findings in cattle [10, 18, 26, 30].

Pinpointing causal mutations is often difficult as many sequence variants may be compatible with the presumed pattern of inheritance (e.g., [18, 24]). In 54 sequenced Braunvieh animals of our study, two intergenic variants were in complete LD with rs383232842. Yet we consider rs383232842 as the most likely to be causal because it results in a substitution of a conserved amino acid residue. Moreover, all compatible noncoding variants are far away from annotated transcripts and are thus less likely to be functionally important. However, it cannot be ruled out *a priori* that such variants affect gene regulation, *e.g.,* if located in a distant enhancer region. A criterion for causality is often that the segregation of the candidate variant is restricted to the population affected by the condition (e.g., [13, 14, 29, 31]). Strictly applying this criterion to our data would have led to the exclusion of rs383232842 as a plausible causal mutation for BH2, since it segregates in Fleckvieh cattle where a previous scan for homozygous haplotype deficiency did not detect homozygosity depletion at the BH2 region [14]. In the present study, we specifically tested the BH2_F_v haplotype with the rs383232842 C-allele for association with juvenile mortality in more than 37,000 Fleckvieh animals. Homozygosity depletion at BH2_FV_ was not significant (P=0.09) possibly because most Fleckvieh animals of the present study were genotyped shortly after birth, before typical signs of disease may become manifest. However, the first-year mortality of descendants is significantly higher in BH2_FV_ risk than non-risk matings (P=2.2x10^−15^) corroborating a detrimental effect of BH2_FV_.

The exchange of genetic material between breeds may result in the manifestation of the same recessively inherited conditions in different populations [27]. Kumar Kadri et al. identified a sequence variant that compromises fertility in three admixed Nordic dairy breeds [25]. Drögemüller et al. mapped a sequence variant for an autosomal recessive skin disease in two geographically separated albeit genetically related dog populations [31]. A diagnostic haplotype associated with progressive degenerative myeloencephalopathy (Weaver syndrome) in Braunvieh cattle also segregates in the Holstein cattle breed [32]. However, the most likely causal variant for Weaver syndrome was never found in haplotype carriers from the Holstein cattle breed indicating that an ancestral version of the haplotype without the Weaver mutation persists in Holstein cattle [32, 33]. The discovery of a recessive mutation that compromises juvenile survival in Fleckvieh and Braunvieh cattle demonstrates for the first time, that deleterious alleles may segregate across closed cattle populations, *e.g.,* populations without recent admixture. Fleckvieh and Braunvieh animals with the rs383232842 C-allele share a common haplotype, indicating they inherited the corresponding chromosome segment from a common ancestor. However, our data neither revealed common ancestors nor provided evidence of an exchange of genetic material between both breeds. A version of the ancestral haplotype without the rs383232842 C-allele also segregates in Holstein cattle at relatively high frequency (5.5 %). These findings indicate that the rs383232842 mutation occurred in an ancient ancestor of Braunvieh and Fleckvieh cattle, however, after the divergence from Holstein cattle, which is compatible with the history of those breeds [34]. While BH2 and BH2_FV_ affect juvenile survival in Braunvieh and Fleckvieh cattle, respectively, the haplotype version without the rs383232842 C-allele does not exhibit homozygous haplotype deficiency in Holstein cattle supporting causality of the rs383232842 polymorphism.

To investigate the pathophysiology that may result from the rs383232842 mutation in *TUBD1,* we examined six homozygous animals. Although the age of the examined animals differed, the disease manifestation was homogeneous: recurrent airway disease resulted in a steadily declining general condition and poor growth performance despite excellent husbandry conditions and medical treatment at the animal clinic. Thus, a more severe disease progression in homozygous animals kept under normal husbandry conditions in conventional farms is likely. The respiratory manifestations resemble disease patterns arising from defective cilia of the respiratory tract [35–37]. We observed microtubular assembly defects in 20-30 % of the cilia of the respiratory epithelium in affected animals which is 4-6 times higher than in healthy individuals [38–40]. Ultrastructural defects of the cilia might also result from infectious or inflammatory diseases [41]. Persistent coughing, tachypnea and airway mucus obstruction in homozygous animals might indicate an impaired mucociliary clearance due to an aberrant ciliary beat pattern [42–45]. We used a cytology brush to collect viable epithelial cells from the nasal mucosa of affected calves to investigate ciliary beat pattern and beat frequency (data not shown). However, all collected samples were excessively covered with mucous exudate and contained numerous inflammatory cells, precluding the analysis of ciliary beating using standard protocols.

Several sequence variants associated with ultrastructural aberrations of respiratory cilia cause chronic airway disease in humans, mice and dogs (e.g., [37, 46, 47]). Physiological respiratory cilia are characterized by a “9x2+2” architecture consisting of a central pair of microtubule surrounded by nine outer microtubule doublets. The basal body is located at the base of the cilium where it attaches to the cell body. Tubulin delta 1 is required for proper microtubule polymerization in basal bodies [48–51]. A lack of TUBD1 causes ultrastructural microtubular defects in basal bodies in *Chlamydomonas* and *Paramecium* [48, 50, 51]. We also observed disorganized microtubules in airway cilia of animals homozygous for the rs383232842 mutation corroborating a crucial role of TUBD1 for proper assembly of cilia. To our knowledge, this is the first report of a phenotypic effect associated with genomic variation in *TUBD1* in a mammalian species. A conserved histidine residue at amino acid position 210 in TUBD1 is involved in microtubule polymerization [49]. Since the rs383232842 C-allele introduces a putatively damaging substitution of the histidine by an arginine, proper microtubule polymerization might be compromised in homozygous animals, resulting in ultrastructural defects of the airway cilia. Besides being involved in cilia assembly, microtubules constitute the bipolar spindle that binds and moves the chromosomes during the different meiotic phases. Disturbed microtubule formation may lead to erroneous segregation of chromosomes with negative effects on growth and differentiation (e.g., [52]). Such disturbances could be responsible for frequently observed prenatal growth retardation and stillbirth of fetuses from BH2 risk matings.

Most homozygous calves (75 %) were stillborn or died shortly after birth. Six hospitalized calves suffered from airway disease early in life. The general condition of all hospitalized animals declined steadily because the disease was not responsive to medical treatment. However, homozygosity for the rs383232842 mutation does not necessarily have fatal consequences as evidenced by a bull homozygous for the putative disease allele without any apparent signs of disease. Although minimizing exposure to environmental pollutants and respiratory pathogens may contribute to a less severe disease progression [53], our results suggest that the mutation is highly penetrant because 17 out of 18 homozygous animals were stillborn or diseased shortly after birth. In any case, our findings now enable to implement efficient genome-based mating strategies to avoid the mating of carrier animals thereby preventing the birth of homozygous calves that are likely to suffer from chronic disease.

## Conclusions

A missense mutation in *TUBD1* is associated with high juvenile mortality in Braunvieh and, unexpectedly, Fleckvieh cattle. Since genomic analyses revealed no evidence for an exchange of genetic material between both breeds, we demonstrate for the first time that the same harmful recessive mutation may segregate in different closed populations, indicating that such mutations may have occurred many generations ago. Homozygous animals suffer from chronic airway disease possibly resulting from defective cilia in the respiratory tract. Our findings now enable the implementation of genome-based mating strategies in order to prevent animal suffering and economic losses.

## Methods

### Animal ethics statement

Homozygous animals were descendants from matings between carrier animals that happened inadvertently in the Braunvieh and Fleckvieh populations prior to the declaration of BH2 carrier bulls. Ear tissue samples of homozygous calves were collected from breeding consultants. Seven homozygous animals were hospitalized and examined at the Clinic for Ruminants of the Veterinary University of Vienna, Austria and at the Clinic of Reproductive Medicine, Vetsuisse-Faculty, University Zurich, Switzerland as part of their regular practice. All hospitalized animals were euthanized because of steadily declining health condition with no prospect of improvement. No ethical approval was required for this study.

### Animals, genotypes, quality control and phasing

Genotypes of 8501 Braunvieh and 37,314 Fleckvieh animals were extracted from the Austrian and German bovine genome databases. Genotypes of 1765 Braunvieh bulls were provided by breeding organisations from Switzerland (n=993), Italy (n=465) and USA (n=316) based on bilateral genotype exchange agreements. All animals had been genotyped with the Illumina BovineSNP50 bead chip (version 1 and version 2) comprising approximately 54,000 SNPs. The chromosomal position of the SNPs was based on the UMD 3.1 assembly of the bovine genome [54]. Mitochondrial, X-chromosomal, Y-chromosomal SNPs and SNPs that had no known chromosomal position were not considered for further analyses. Animals with a call-rate less than 95 % were excluded as well as animals with mendelian conflicts with their genotyped sires at more than 200 SNPs. We excluded SNPs with a minor allele frequency below 0.5 % as well as SNPs with call rate less than 90 % or more than 0.5 % mendelian conflicts in sire-offspring pairs. The final Braunvieh data consisted of 8,446 animals and 35,382 autosomal SNPs with an average per-individual call-rate of 99.61 %. The final Fleckvieh data consisted of 37,314 animals and 41,027 autosomal SNPs with an average per-individual call-rate of 99.82%. Haplotypes for BTA19 were inferred and sporadically missing genotypes were imputed using the *shapeit* software (v2.778) [55].

### Mapping of BH2 in the Braunvieh cattle population

A sliding window with variable size (ranging from 0.75 to 8 Mb) was shifted along chromosome 19 (in steps of 0.5 x window size). Within each sliding window, the observed number of homozygotes was compared to the expected number for all haplotypes with a frequency above 3% using Fisher exact tests. The expected number of homozygous animals was calculated using haplotype information from sire, maternal grandsire and haplotype frequency. Phenotypic effects were analyzed for all haplotypes that showed significant depletion of homozygous animals (P<1 x 10^−4^).

### Mapping of BH2FV in the Fleckvieh cattle population

To identify a haplotype specific for the rs383232842 C-allele, we exploited haplotypes of 10,363 Fleckvieh animals that had been genotyped and (partially) imputed at 652,853 SNPs [24]. We extracted haplotypes of 1947 animals that had also been genotyped at rs383232842 (see below). A haplotype window consisting of five adjacent SNPs was centered at rs383232842 and increased by one SNP on either side as long as a common haplotype (BH2_FV_) was heterozygous in all animals that were heterozygous at rs383232842. To assess the frequency of BH2_FV_ in a representative sample of the Fleckvieh cattle population, we analysed haplotypes of 37,314 Fleckvieh animals from the bovine genome databases (see above).

### Mapping of BH2 in the Holstein cattle population

To test if the common BH2 / BH2_FV_ haplotype also segregates in the Holstein cattle breed, we used haplotypes of 8840 Holstein artificial insemination bulls that had been genotyped previously with the Illumina BovineSNP50 Bead chip [56]. BH2 carriers were identified based on eight SNPs (from 10,694,269 bp to 11,435,269 bp) that were common in BH2 / BH2FV carriers, respectively.

### Phenotypic effects associated with BH2 and BH2_FV_

Insemination success was analyzed based on 55,837 artificial inseminations that were carried out with Braunvieh bulls that had a known haplotype status for BH2. The insemination success in risk matings (carrier bulls mated to daughters of carrier sires) was compared with non-risk matings (non-carrier bulls mated to daughters of carrier sires) using a logistic regression model. Perinatal and first-year mortality was analyzed in 32,100 and 46,566 Braunvieh and Fleckvieh calvings, respectively, with known haplotype states from sires and maternal grandsires. Calves that left the recording system within 365 days after birth for other reasons than death (e.g., export to other countries) were not considered. A Kaplan-Meier estimator was obtained by comparing the first-year survival rate of calves from risk matings with non-risk matings [57].

### Generation of sequence data

Genomic DNA of 290 animals (149 Fleckvieh, 54 Braunvieh, 51 Holstein, 15 Original Simmental, 12 Gelbvieh, 7 Northern Finncattle, 2 Ayrshire) was prepared from semen and blood samples following standard DNA extraction protocols. Paired-end libraries were prepared using the paired-end TruSeq DNA sample prep kit (Illumina inc., San Diego, CA, USA) and sequenced using the Illumina HiSeq 2500 instrument (Illumina inc., San Diego, CA, USA). The resulting reads were processed with the Illumina BaseCaller during the sequencing step. The alignment of the reads to the University of Maryland reference sequence (UMD3.1) [54] was performed using the *BWA mem* algorithm [58],[59]. The resulting per individual SAM files were converted into BAM files with *SAMtools* [60]. Duplicate reads were identified and marked with the *MarkDuplicates* command of *Picard* [61]. Part of the sequencing data were previously generated [14, 62] and were contributed to the 1000 bull genomes project [21].

### Variant calling and imputation

SNPs, short insertions and deletions were genotyped in 290 sequenced animals simultaneously using the multi-sample approach implemented in *mpileup* of *SAMtools* along with *BCFtools* [60]. *Beagle* [63] phasing and imputation was used to improve the primary genotype calling by *SAMtools*. The detection of structural variants was performed in 227 sequenced animals with an average genome fold coverage higher than 10x using the *Pindel* software package [64]. The functional effects of polymorphic sites were predicted based on the annotation of the UMD3.1 assembly of the bovine genome using the *Variant Effect Predictor* tool from Ensembl [65],[66]. The consequences of non-synonymous amino acid substitutions on protein function were predicted using *PolyPhen-2* [67] and *SIFT* [68].

### Identification of BH2-associated variants

We screened 110,294 polymorphic sites within a 9.71 Mb interval (between 3.67 Mb and 13.38 Mb) on BTA19 encompassing BH2 for variants that were homozygous for the alternate allele in BH2_hom_, heterozygous in five BH2 carriers and homozygous for the reference allele in 48 control animals of the Braunvieh population. The genotype distributions of three variants that fulfilled these criteria were assessed in 236 sequenced animals representing six cattle breeds other than Braunvieh.

### Validation of the rs383232842 polymorphism

Genotypes for rs383232842 were obtained in 661 Braunvieh, 3807 Fleckvieh and 503 Holstein animals using a customized KASP^TM^ genotyping assay (LGC Genomics) (FAM/HEX/reverse primer sequence:
TGTTCATGAGAACGATGCTGTTCG/CTTGTTCATGAGAACGATGCTGTTCA/TGCTTGATATTCATCAGCTTCACACAGAT).

### Clinical examinations

Seven animals were hospitalized for a period of 19 to 164 days at the animal clinic. Initial examination (including weighing) was performed upon admission. Weight records and blood samples from three animals were collected once or twice a week. Red and white blood cell count was determined using an automatic hematology analyzer (ADVIA 2120i, Siemens Healthcare, Vienna, Austria). Glucose, total protein, albumin, cholesterol, NEFA, total bilirubin, creatinine, potassium, sodium, AST, GLDH, LDH were measured in blood plasma using an automatic analyzer (Cobas 6000/c501, Roche Diagnostics GmbH, Rotkreuz, Switzerland).

### Pathological examinations

All hospitalized animals were euthanized at 32 to 502 days of age because they suffered from chronic disease with no prospect of improvement. Necropsy and collection of tissue samples were done immediately after euthanasia. For histology, samples from nasal mucosa, trachea, lung, heart, liver, kidney, pancreas, small and large intestine, mesenteric lymph node, intestine were fixed in 10 % buffered formalin and embedded in paraffin wax. Sections (3 μm) were stained with hematoxylin and eosin (HE) and alcian blue. For transmission electron microscopy, samples from nasal mucosa, trachea, bronchi and bronchioles were collected from four animals and fixed in 5 % glutaraldehyde (Merck, Darmstadt, Germany) in 0.1 M phosphate buffer (Sigma-Aldrich, Vienna, Austria), pH 7.2, at 4°C for 3 h and then postfixed in 1 % osmium tetroxide (Merck, Darmstadt, Germany) in the same buffer at 4°C for 2 h. After dehydration in an alcohol gradient series and propylene oxide (Merck, Darmstadt, Germany), the tissue samples were embedded in glycidyl ether 100 (Serva, Heidelberg, Germany). The ultrathin sections were cut on a Leica Ultramicrotome (Leica Ultracut S, Vienna, Austria) and stained with uranyl acetate (Sigma-Aldrich, Vienna, Austria) and lead citrate (Merck, Darmstadt, Germany). Ultrathin sections were examined with a Zeiss TEM 900 electron microscope (Carl Zeiss, Oberkochen, Germany) operated at 50 kV.

## Availability of supporting data

Whole-genome sequencing data of BH2_hom_ were deposited in the European Nucleotide Archive (http://www.ebi.ac.uk/ena) under accession number PRJEB12807.

## List of abbreviations

BH2: Braunvieh haplotype 2;
BV: Braunvieh;
FV: Fleckvieh;
LD: linkage disequilibrium;
SNP: Single nucleotide polymorphism;
TUBD1: Tubulin delta 1;

## Authors’ contribution

HP, HS and RF conceived the study, designed the experiments and analyzed the data; HS and CF analyzed genotype and phenotype data; HP analyzed sequence and genotype data; HS, FS, RW and OG conceived the calf monitoring project; JB, TW, ND, MH, HW and UB examined homozygous animals; CW carried out the sequencing experiments, CW and SJ carried out the molecular genetic experiments, BFW and MD contributed to the design of the experiment, HP wrote the manuscript.

## Acknowledgements

We acknowledge support from Braunvieh Austria and Braunvieh Schweiz during sample collection. We acknowledge the InterGenomics partners (Arbeitsgemeinschaft der Österreichischen BraunviehZüchter, Austria; Arbeitsgemeinschaft Deutsches Braunvieh, Germany; Associazione nazionale allevatori bovini della razza Bruna, Italy; Brown Swiss Association of the US, USA; Brune Genetique Services, France; Braunvieh Schweiz, Switzerland; and Zveza rejcev govedi rjave pasme Slovenije, Slovenia) for contribution of genotypes to this study. We thank the Arbeitsgemeinschaft Süddeutscher Rinderzüchter und Besamungsorganisationen e.V. (ASR), the Arbeitsgemeinschaft österreichischer Fleckviehzüchter (AGÖF) and the Förderverein Biologietechnologieforschung e.V. (FBF) for providing genotype data. We thank Associazione Nazionle Allevatori Bovini della Razza Bruna Italiana (ANARBI) and Arbeitsgemeinschaft Deutsches Braunvieh for providing pictures of affected animals and support in sample collection.

## Additional files

### Additional file 1 (Additional_file_1.xlsx)

Title: **Genotype distribution of 52 sequence variants in LD with BH2.**

Description: Non-reference allele frequency and genotype distribution of 52 variants in LD with BH2 in 290 animals representing nine cohorts (number of animals homozygous for the reference allele | number of heterozygous animals | number of animals homozygous for the non-reference allele). Grey background highlights a synonymous variant in *PPM1E*.). Bold type indicates five variants that were located within the 1.14 Mb BH2 haplotype. Blue background indicates three variants that were not homozygous among 1682 animals that had been sequenced in the 1000 bull genomes project. Red color highlights a missense variant (rs383232842) in the *TUBD1* gene.

### Additional file 2 (Additional_file_2.txt)

Title: **Functional effects of 52 variants in LD with BH2.**

Description: The functional consequences of the non-reference alleles were predicted using the *Variant Effect Predictor* tool from Ensembl.

### Additional file 3 (Additional_file_3.pdf)

Title: **Three variants compatible with recessive inheritance of BH2.**

Description: Genotypes of three sequence variants compatible with recessive inheritance of BH2 in 290 sequenced animals representing seven cattle breeds. The sequenced Braunvieh animals were classified into carrier and non-carrier animals using haplotypes inferred from array-derived genotypes. Blue color indicates the missense mutation in *TUBD1*.

### Additional file 4 (Additional_file_4.pdf)

Title: **Across-species conservation of tubulin delta 1.**

Description: Part of the multispecies alignment of the TUBD1 protein sequence. Blue color highlights the missense mutation.

### Additional file 5 (Additional_file_5.xlsx)

Title: **Genotype distribution of three variants in LD with BH2.**

### Additional file 6 (Additional_file_6.xlsx)

Title: **Haplotype analysis in Braunvieh, Fleckvieh and Holstein cattle.**

Description: The BH2 and BH2_FV_ haplotypes were detected using 777K and 50K genotyping data, respectively. Yellow and red color highlights eight common SNPs in Braunvieh and Fleckvieh cattle and the missense variant in *TUBD1*, respectively. Note that the BH2 haplotype version in Holstein cattle does not contain the rs383232842 C-allele.

### Additional file 7 (Additional_file_7.tif)

Title: **Cluster analyses in 280 Braunvieh and 442 Fleckvieh animals.**

Description: Principal component analysis (PCA) using genotypes of 42,056 autosomal SNPs (a) and 1121 SNPs located on chromosome 19 (b), respectively. All animals were born between 1970 and 2010. The PCA separated the animals by breed without any evidence for an admixture. There was no indication that BH2 and BH2FV carriers were more closely related to each other than to non-carrier animals. Hierarchical clustering using genotypes of 1121 SNPs located on chromosome 19 (c). Animals were clearly separated by breed without any evidence of an admixture of BH2 and BH2_FV_ carriers.

### Additional file 8 (Additional_file_8.png)

Title: **Photograph of a stillborn homozygous calf.**

Description: Despite a normal gestation length (283 days), the calf was underweight at birth at 25 kg. The head of the stillborn calf appears elongated and the limbs were thin and elongated. The analysis of histological sections of a diverse tissue panel revealed no pathological findings.

### Additional file 9 (Additional_file_9.tif)

Title: **Growth of six BH2 homozygous calves.**

Description: The weight of six BH2 homozygous calves (BV1-5, FV1) at admission to the clinic and at euthanasia is compared to 74,422 healthy Fleckvieh animals (grey boxes) and three Fleckvieh animals with loss of function (LoF) variants in *GON4L* and *SLC2A2* that manifest in growth retardation [22],[14].

### Additional file 10 (Additional_file_l0.tif)

Title: **Blood parameters of BV1, BV2 and FV1.**

Description: The grey shaded areas represent reference values that were determined based on [69].

### Additional file 11 (Additional_file_ll.png)

Title: **Macroscopic lesions in the respiratory tract in four homozygous animals.**

Description: Congestion and mucopurulent inflammation of the nasal conchae of BV2 (a). Different grades of obstruction, atelectasis and/or bronchopneumonia (b-d): single consolidated area in the right cranial lung lobe of BV1 (b), multiple lobular consolidations in both cranial and middle lung lobes of FV1 (c) and severe pneumonic induration of the majority of both cranial and middle lung lobes of BV5 (d). Arrows indicate the border to physiological lung tissue.

### Additional file 12 (Additional_file_12.png)

Title: **Histological lesions in the respiratory tract of two homozygous animals.**

Description: The ciliated epithelium of the nasal mucosa and the underlying submucosa of FV1 are infiltrated with inflammatory cells, particularly neutrophils (a). Lung lesions of BV5 including hyperplastic bronchiolar epithelium, partial obstruction of the bronchiolar lumen by purulent exsudate and consolidation of the surrounding lung tissue due to obstructive atelectasis and mild infiltration with neutrophils (b).

### Additional file 13 (Additional_file_13.tif)

Title: **Three animals homozygous for the rs383232842 C-allele.**

Description: A homozygous young bull (BV6) at the age of 370 days (a). According to the possessing farmer, the bull did not suffer from respiratory disease in the past. The weight of another homozygous young bull (BV7) at the age of 271 days was 243 kg (b). The possessing farmer reported that the young bull suffered from life-threatening bronchopneumonia early in life. Although the animal recovered from the disease, coughing and excessive mucous exudation from the nostrils were apparent during its entire life. Multiple lung lesions were noticed after slaughter. A female calf homozygous (BV8) for the rs383232842 C-allele at the age of 194 days with an unaffected coeval (c-e). Photographs were kindly provided by Franz Birkenmaier (a, c-e) and Attilio Rossoni (b).

